# Genome sequence of *Bacillus paranthracis* strain SCM10-01, isolated from the intestinal mucosa of a wild *Synallaxis cabanisi* collected in Peru

**DOI:** 10.64898/2026.08.12.744436

**Authors:** Eric Finkelstein, Sarah M. Hird

## Abstract

We report the genome sequence of *Bacillus paranthracis* SCM10-01, isolated from a wild neotropical bird (*Synallaxis cabanisi*) collected in Peru. The assembly yielded one chromosome, three plasmids, and Bacillus phage SCM10. Genomic screening identified complete hemolysin BL, nonhemolytic enterotoxin operons, and cytotoxin K2, but no anthrax-associated toxin or capsule genes.

## Announcement

The *Bacillus cereus* group comprises taxonomically ambiguous bacteria of medical, veterinary, and agricultural importance (1, 2). Here, we report the genome of *Bacillus paranthracis* SCM10-01 and assess its taxonomy and virulence genes.

A *Synallaxis cabanisi* (Cabanis’s Spinetail) was collected in Peru in 2016 (LSUMNS B88879); the intestine stored at -80°C. In 2023, the tissue was opened and the contents were removed. It was rinsed with phosphate-buffered saline, suspended in phosphate freezing buffer and sonicated for 10 min in a Branson M5800 water bath (3). The sonicate was enriched in tryptic soy broth at 36°C for 24 h with shaking, serially diluted 10-fold, plated on tryptic soy agar, incubated at 40°C for 24 h, and single-colony purified. Genomic DNA was extracted using the Qiagen QIAamp PowerFecal Pro DNA Kit.

Library preparation and sequencing were conducted at the University of Connecticut Center for Genome Innovation. Illumina libraries were prepared using the Nextera XT kit and sequenced on a MiSeq platform, 2x250 bp paired-end. Nanopore libraries were prepared using Rapid Barcoding Kit (SQK-RBK114.24) with SPRI size selection and protocol modifications (4). Sequencing was performed on a GridION using an R10.4.1 flow cell and super-accuracy base called in MinKNOW v25.09.18.

Read filtering, subsampling, hybrid-assembly, and polishing were performed using Hybracter v0.12.0 (5) (hybrid-single; Flye --nano-hq --auto; Polypolish and Pypolca --careful). Nanopore read metrics were calculated with NanoPlot v1.46.2 (6). Replicons were screened for redundancy using MUMmer v4.0.2 (7) (nucmer --maxmatch; show-coords --rclT). Assembly adaptor and vector contamination were screened and removed using NCBI FCS-adaptor v0.5.5 (8) (--prok). Assembly quality was assessed by read mapping (BWA-MEM2 v2.2.1 (9), Minimap2 v2.28 (10); SAMtools v1.21 (11) stats/coverage), QUAST v5.3.0 (12), and BUSCO v6.0.0 (13) (bacillus_odb12). Proviruses and viral replicons were identified with geNomad v1.12.0 (14) (database v1.9) and CheckV v1.1.0 (15) (database v1.5).

Taxonomy, annotation, and virulence genes were determined with BTyper3 v3.4.0 (16), Bakta v1.11.4 (17) (2025.02.24 database), and AMRFinderPlus v4.2.7 (18) (2026-05-15.1 database) respectively. Type strains sharing ≥93% average nucleotide identity (ANI) with SCM10-01 were retrieved with NCBI Datasets v2 (19). Pairwise ANI was calculated with Skani v0.3.1 triangle (20) and clustered by UPGMA in SciPy v1.15.3 (21).

The hybrid assembly produced one contiguous chromosome and four circular replicons (Table 1). Illumina and Nanopore reads were mapped to 99.99% and 99.3% respectively. A 1,016 bp linear contig was excluded as a redundant assembly fragment because >90% of its length aligned to the existing assembly sequences, primarily to the 211 kb plasmid. geNomad classified one plasmid as viral, which CheckV scored as high-quality and near complete (99.3% completion, 0% contamination); two additional proviruses were predicted, one chromosomal and one plasmid-associated.

**Table 1.**
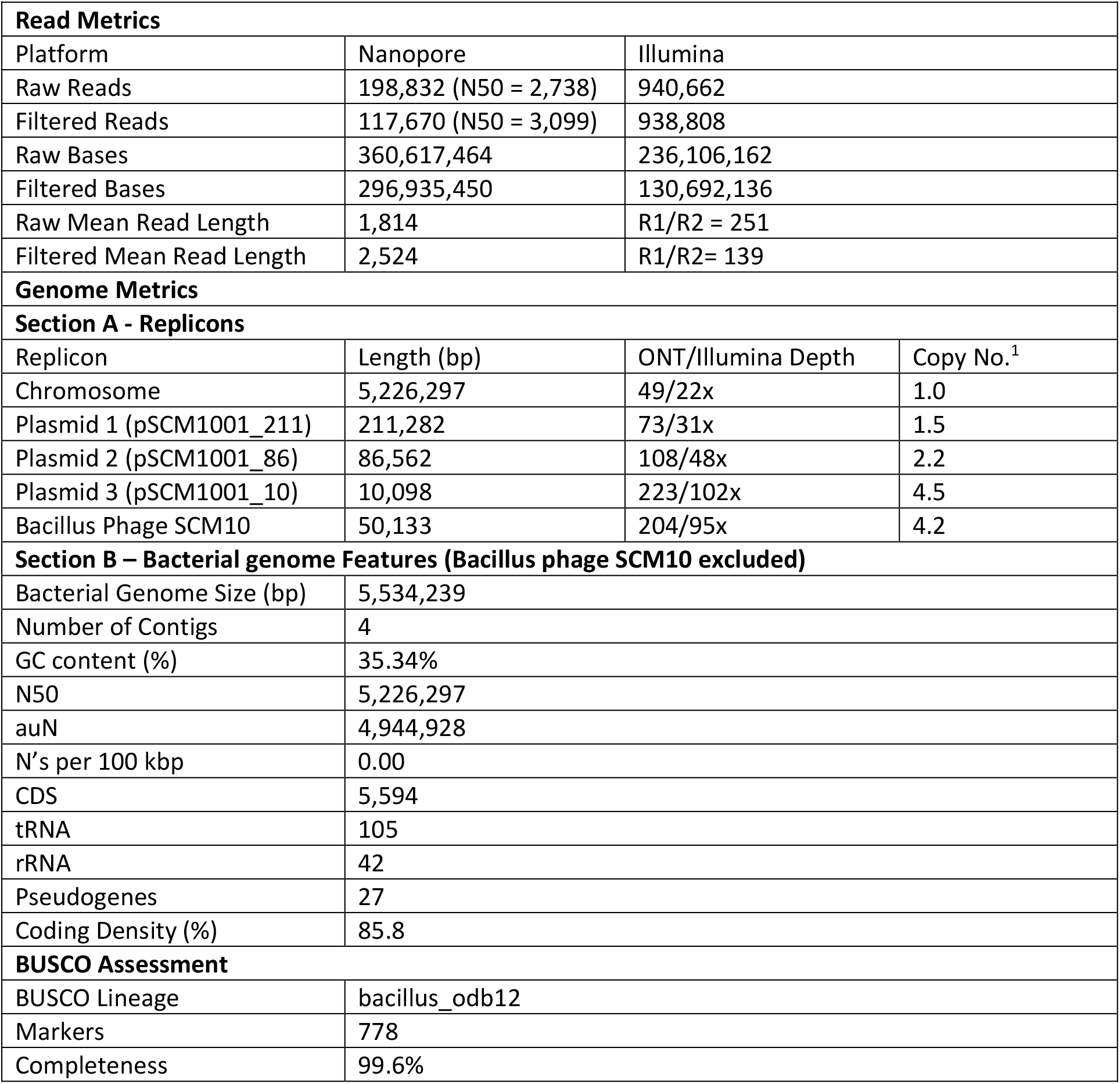
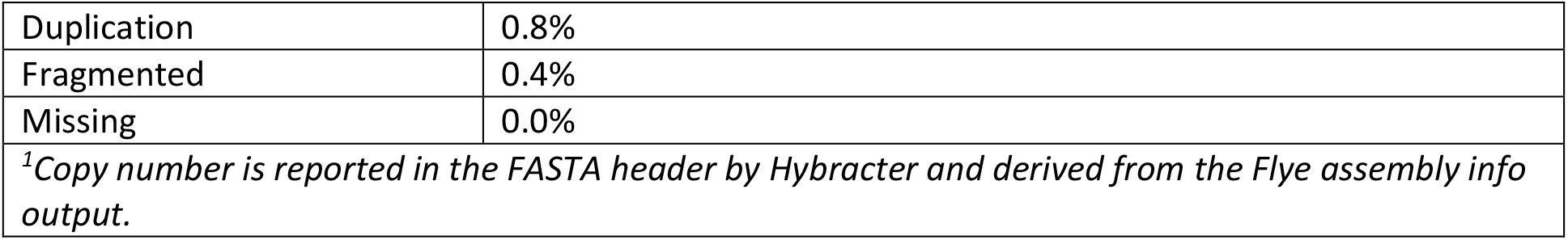
Sequencing, Assembly, and Genome Metrics.

| Table 1. Sequencing, Assembly, and Genome Metrics |  |  |  |
| --- | --- | --- | --- |
| Read Metrics |  |  |  |
| Platform | Nanopore | Illumina |  |
| Raw Reads | 198,832 (N50 = 2,738) | 940,662 |  |
| Filtered Reads | 117,670 (N50 = 3,099) | 938,808 |  |
| Raw Bases | 360,617,464 | 236,106,162 |  |
| Filtered Bases | 296,935,450 | 130,692,136 |  |
| Raw Mean Read Length | 1,814 | R1/R2 = 251 |  |
| Filtered Mean Read Length | 2,524 | R1/R2= 139 |  |
| Genome Metrics |  |  |  |
| Section A - Replicons |  |  |  |
| Replicon | Length (bp) | ONT/Illumina Depth | Copy No.<sup>1</sup> |
| Chromosome | 5,226,297 | 49/22x | 1.0 |
| Plasmid 1 (pSCM1001_211) | 211,282 | 73/31x | 1.5 |
| Plasmid 2 (pSCM1001_86) | 86,562 | 108/48x | 2.2 |
| Plasmid 3 (pSCM1001_10) | 10,098 | 223/102x | 4.5 |
| Bacillus Phage SCM10 | 50,133 | 204/95x | 4.2 |
| Section B – Bacterial genome Features (Bacillus phage SCM10 excluded) |  |  |  |
| Bacterial Genome Size (bp) | 5,534,239 |  |  |
| Number of Contigs | 4 |  |  |
| GC content (%) | 35.34% |  |  |
| N50 | 5,226,297 |  |  |
| auN | 4,944,928 |  |  |
| N's per 100 kbp | 0.00 |  |  |
| CDS | 5,594 |  |  |
| tRNA | 105 |  |  |
| rRNA | 42 |  |  |
| Pseudogenes | 27 |  |  |
| Coding Density (%) | 85.8 |  |  |
| BUSCO Assessment |  |  |  |
| BUSCO Lineage | bacillus_odb12 |  |  |
| Markers | 778 |  |  |
| Completeness | 99.6% |  |  |

|  |  |
| --- | --- |
| Duplication | 0.8% |
| Fragmented | 0.4% |
| Missing | 0.0% |
| <sup>1</sup>Copy number is reported in the FASTA header by Hybracter and derived from the Flye assembly info output. |  |

BTyper3 assigned SCM10-01 to *Bacillus mosaicus* genomospecies without biovar, serovar, or subspecies designation. SCM10-01 showed the highest ANI with type strains *Bacillus anthracis* Vollum (97.75%) and Bacillus paranthracis Mn5 (95.34%). AMRFinderPlus predicted an intact *plcR*, complete *hblCDAB* and *nheABC* operons, and *cytK2* in SCM10-01, but did not detect anthrax-associated virulence markers *atxA, pagA, lef, cya*, or *capBCADE* (Fig. 1). Given the overlapping species boundaries within *Bacillus mosaicus* and its absence from NCBI taxonomy, SCM10-01 is conservatively reported under the deposited designation *Bacillus paranthracis*.

**Figure 1.**
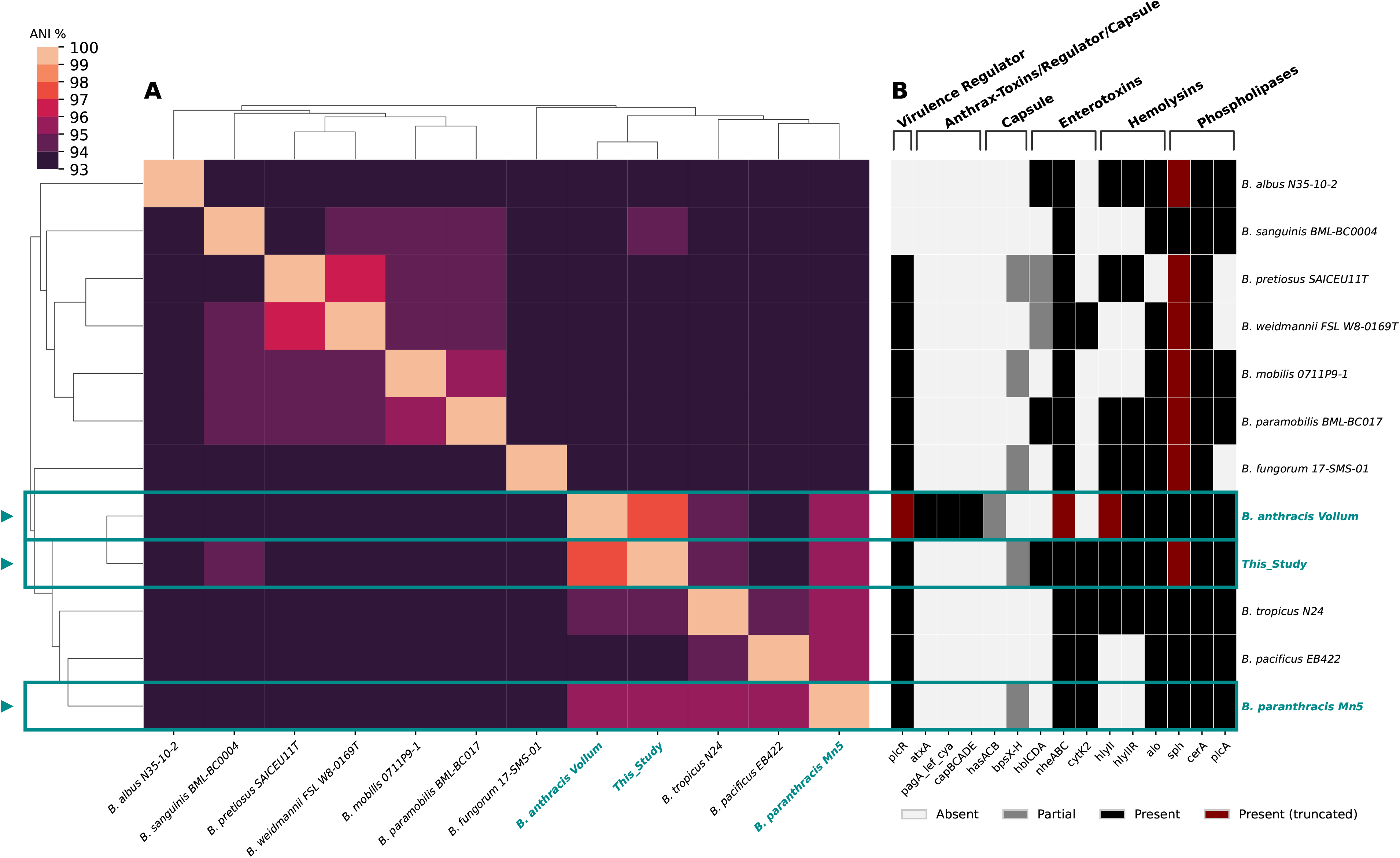
Genome similarity and selected virulence-marker profiles of *Bacillus paranthracis* SCM10-01 and 11 related *Bacillus cereus* group type strains.

Type strains sharing ≥93% ANI with SCM10-01, as identified by BTyper3, were included. (A) Pairwise ANI values were calculated with Skani triangle, converted to distances of 100 – ANI, and hierarchically clustered with UPGMA using average linkage. The ANI heatmap is displayed with a lower scale limit of 93%. (B) Selected virulence marker profiles were determined with AMRFinderPlus and are shown in the same genome order as in (A). Markers represent a curated, nonexhaustive set relevant to *Bacillus cereus* group pathogenesis and anthrax-associated virulence and are grouped by functional category, indicated by brackets. Cell colors denote the marker status as absent, partial, present, or present with an AMRFinderPlus predicted internal stop codon. *Bacillus paranthracis* SCM10-01 (this study) and its two closest reference strain, *Bacillus anthracis* Vollum and *Bacillus paranthracis* Mn5 are highlighted in blue. The emetic toxin *ces* operon and insecticidal *cry, cyt, vip*, and *sip* genes were omitted because they were not detected in any of the genomes.

## Data availability statement

The genome sequence of *Bacillus paranthracis* SCM10-01 is processing in NCBI WGS under accession number JCAQAB000000000. Raw Illumina and Oxford Nanopore reads are available in SRA under accession numbers SRX33992432 and SRX33992433 respectively (BioProject accession no. PRJNA1480000 and BioSample accession no. SAMN61006589). The Bacillus phage SCM10 was deposited under a separate BioProject PRJNA1490058 (accession number [PZ686256](https://www.ncbi.nlm.nih.gov/nuccore/3369854480)), and shares the same BioSample as strain SCM10-01. The JupyterLab notebook and corresponding data used to generate Figure 1 is available through figshare 10.6084/m9.figshare.33111947.

## Acknowledgments

We thank Ryan Duggan and Phoenix Wang for isolation and cultivation of strain SCM10-01 and genomic DNA extraction; The University of Connecticut Center for Genome Innovation for Illumina library preparation and sequencing; and Dr. Kendra Maas for assistance with Oxford Nanopore library preparation and sequencing. We thank the LSU Museum of Natural Science Collection of Genetic Resources for the tissue loan. This study was supported by the National Science Foundation (award no. 2030460).

## Notes

https://www.ncbi.nlm.nih.gov/sra/?term=SRX33992432

https://www.ncbi.nlm.nih.gov/sra/?term=SRX33992433

https://www.ncbi.nlm.nih.gov/bioproject/?term=PRJNA1480000

https://www.ncbi.nlm.nih.gov/biosample/?term=SAMN61006589

https://figshare.com/search?q=10.6084%2Fm9.figshare.33111947

### Competing Interest Statement

The authors have declared no competing interest.

## References

1. Okamoto A, Okutani A. 2024. The Bacillus cereus group, p. 957–986. In Molecular Medical Microbiology. Elsevier.

2. Carroll LM, Cheng RA, Wiedmann M, Kovac J. 2022. Keeping up with the Bacillus cereus group: taxonomy through the genomics era and beyond. Critical Reviews in Food Science and Nutrition 62:7677–7702.

3. Morella NM, Weng FC-H, Joubert PM, Metcalf CJE, Lindow S, Koskella B. 2020. Successive passaging of a plant-associated microbiome reveals robust habitat and host genotype-dependent selection. Proc Natl Acad Sci USA 117:1148–1159.

4. Finkelstein E. 2026. Whole-Genome Nanopore Sequencing of Bacterial gDNA (v1). protocols.io.

5. Bouras G, Houtak G, Wick RR, Mallawaarachchi V, Roach MJ, Papudeshi B, Judd LM, Sheppard AE, Edwards RA, Vreugde S. 2024. Hybracter: enabling scalable, automated, complete and accurate bacterial genome assemblies. Microbial Genomics 10.

6. De Coster W, Rademakers R. 2023. NanoPack2: population-scale evaluation of long-read sequencing data. Bioinformatics 39:btad311.

7. Marçais G, Delcher AL, Phillippy AM, Coston R, Salzberg SL, Zimin A. 2018. MUMmer4: A fast and versatile genome alignment system. PLoS Comput Biol 14:e1005944.

8. Astashyn A, Tvedte ES, Sweeney D, Sapojnikov V, Bouk N, Joukov V, Mozes E, Strope PK, Sylla PM, Wagner L, Bidwell SL, Brown LC, Clark K, Davis EW, Smith-White B, Hlavina W, Pruitt KD, Schneider VA, Murphy TD. 2024. Rapid and sensitive detection of genome contamination at scale with FCS-GX. Genome Biol 25:60.

9. Vasimuddin Md, Misra S, Li H, Aluru S. 2019. Efficient Architecture-Aware Acceleration of BWA-MEM for Multicore Systems, p. 314–324. In 2019 IEEE International Parallel and Distributed Processing Symposium (IPDPS). IEEE, Rio de Janeiro, Brazil.

10. Li H. 2018. Minimap2: pairwise alignment for nucleotide sequences. Bioinformatics 34:3094–3100.

11. Danecek P, Bonfield JK, Liddle J, Marshall J, Ohan V, Pollard MO, Whitwham A, Keane T, McCarthy SA, Davies RM, Li H. 2021. Twelve years of SAMtools and BCFtools. GigaScience 10:giab008.

12. Gurevich A, Saveliev V, Vyahhi N, Tesler G. 2013. QUAST: quality assessment tool for genome assemblies. Bioinformatics 29:1072–1075.

13. Simão FA, Waterhouse RM, Ioannidis P, Kriventseva EV, Zdobnov EM. 2015. BUSCO: assessing genome assembly and annotation completeness with single-copy orthologs. Bioinformatics 31:3210–3212.

14. Camargo AP, Roux S, Schulz F, Babinski M, Xu Y, Hu B, Chain PSG, Nayfach S, Kyrpides NC. 2024. Identification of mobile genetic elements with geNomad. Nat Biotechnol 42:1303–1312.

15. Nayfach S, Camargo AP, Schulz F, Eloe-Fadrosh E, Roux S, Kyrpides NC. 2021. CheckV assesses the quality and completeness of metagenome-assembled viral genomes. Nat Biotechnol 39:578–585.

16. Carroll LM, Cheng RA, Kovac J. 2020. No Assembly Required: Using BTyper3 to Assess the Congruency of a Proposed Taxonomic Framework for the Bacillus cereus Group With Historical Typing Methods. Front Microbiol 11:580691.

17. Schwengers O, Jelonek L, Dieckmann MA, Beyvers S, Blom J, Goesmann A. 2021. Bakta: rapid and standardized annotation of bacterial genomes via alignment-free sequence identification: Find out more about Bakta, the motivation, challenges and applications, here. Microbial Genomics 7.

18. Feldgarden M, Brover V, Gonzalez-Escalona N, Frye JG, Haendiges J, Haft DH, Hoffmann M, Pettengill JB, Prasad AB, Tillman GE, Tyson GH, Klimke W. 2021. AMRFinderPlus and the Reference Gene Catalog facilitate examination of the genomic links among antimicrobial resistance, stress response, and virulence. Sci Rep 11:12728.

19. O’Leary NA, Cox E, Holmes JB, Anderson WR, Falk R, Hem V, Tsuchiya MTN, Schuler GD, Zhang X, Torcivia J, Ketter A, Breen L, Cothran J, Bajwa H, Tinne J, Meric PA, Hlavina W, Schneider VA. 2024. Exploring and retrieving sequence and metadata for species across the tree of life with NCBI Datasets. Sci Data 11:732.

20. Shaw J, Yu YW. 2023. Fast and robust metagenomic sequence comparison through sparse chaining with skani. Nat Methods 20:1661–1665.

21. Virtanen P, Gommers R, Oliphant TE, Haberland M, Reddy T, Cournapeau D, Burovski E, Peterson P, Weckesser W, Bright J, Van Der Walt SJ, Brett M, Wilson J, Millman KJ, Mayorov N, Nelson ARJ, Jones E, Kern R, Larson E, Carey CJ, Polat İ, Feng Y, Moore EW, VanderPlas J, Laxalde D, Perktold J, Cimrman R, Henriksen I, Quintero EA, Harris CR, Archibald AM, Ribeiro AH, Pedregosa F, Van Mulbregt P, SciPy 1.0 Contributors, Vijaykumar A, Bardelli AP, Rothberg A, Hilboll A, Kloeckner A, Scopatz A, Lee A, Rokem A, Woods CN, Fulton C, Masson C, Häggström C, Fitzgerald C, Nicholson DA, Hagen DR, Pasechnik DV, Olivetti E, Martin E, Wieser E, Silva F, Lenders F, Wilhelm F, Young G, Price GA, Ingold G-L, Allen GE, Lee GR, Audren H, Probst I, Dietrich JP, Silterra J, Webber JT, Slavič J, Nothman J, Buchner J, Kulick J, Schönberger JL, De Miranda Cardoso JV, Reimer J, Harrington J, Rodríguez JLC, Nunez-Iglesias J, Kuczynski J, Tritz K, Thoma M, Newville M, Kümmerer M, Bolingbroke M, Tartre M, Pak M, Smith NJ, Nowaczyk N, Shebanov N, Pavlyk O, Brodtkorb PA, Lee P, McGibbon RT, Feldbauer R, Lewis S, Tygier S, Sievert S, Vigna S, Peterson S, More S, Pudlik T, Oshima T, Pingel TJ, Robitaille TP, Spura T, Jones TR, Cera T, Leslie T, Zito T, Krauss T, Upadhyay U, Halchenko YO, Vázquez-Baeza Y. 2020. SciPy 1.0: fundamental algorithms for scientific computing in Python. Nat Methods 17:261–272.

